# E-cadherin mechanotransduction activates EGFR-ERK signaling in epithelial cells by inducing ADAM-mediated ligand shedding

**DOI:** 10.1101/2025.03.06.641828

**Authors:** Ronja M. Houtekamer, Mirjam C. van der Net, Marjolein J. Vliem, Tomas E.J.C. Noordzij, Lisa van Uden, Robert M. van Es, Joo Yong Sim, Eriko Deguchi, Kenta Terai, Matthew A. Hopcroft, Harmjan R. Vos, Beth P. Pruitt, Michiyuki Matsuda, Willem-Jan Pannekoek, Martijn Gloerich

**Author notes:** Corresponding authors. (W.J.P.), (M.G.).

## Abstract

The behavior of cells is governed by signals originating from their local environment, including mechanical forces that cells experience. Forces are transduced by mechanosensitive proteins, which can impinge on signaling cascades that are also activated by growth factor receptors upon ligand binding. We investigated the crosstalk between these mechanical and biochemical signals in the regulation of intracellular signaling networks in epithelial monolayers. Phosphoproteomic and transcriptomic analyses on epithelial monolayers subjected to mechanical strain revealed ERK signaling as a predominant strain-activated hub, initiated at the level of the upstream epidermal growth factor receptor (EGFR). Strain-induced EGFR-ERK signaling depends on mechanosensitive E-cadherin adhesions. Proximity labeling identified the metalloproteinase ADAM17, an enzyme that mediates shedding of soluble EGFR ligands, to be closely associated with E-cadherin. We developed a novel probe for monitoring ADAM-mediated shedding, which demonstrated that mechanical strain induced ADAM activation. Mechanically-induced ADAM activation was essential for mechanosensitive signaling from E-cadherin adhesions towards EGFR-ERK. Collectively, our data demonstrate that mechanical strain transduced by E-cadherin adhesion triggers the shedding of EGFR ligands that stimulate downstream downstream ERK activity. Our findings illustrate how mechanical signals and biochemical ligands can operate within a single, linear signaling cascade.

**Significance statement:** Cells integrate different types of information that they receive from their local environment to regulate their behavior. This includes biochemical signals, such as growth factors binding to their dedicated receptor. Similarly, cells respond to mechanical forces that they are subjected to. Although biochemical and mechanical signals can elicit similar signaling responses in cells, the interplay between these types of signals is not well understood. Here we unveil that mechanical strain of epithelia modulates the activity of the EGFR-ERK signaling pathway by controlling the availability of growth factors that bind and activate EGFR. This finding demonstrates that biochemical and mechanical signals do not act in a segregated fashion, but rather can function in a linear cascade, shedding light on fundamental principles governing cellular regulation.

## Introduction

Epithelia are dynamic tissues in which individual cells adapt their behavior, including the rate of proliferation, to the need of the tissue. Cells integrate these needs for behavioral changes from a myriad of instructive signals received from their local surroundings, including neigboring cells. These extrinsic signals can be of biochemical nature, such as growth factors, but also encompass mechanical forces that cells exert on one another and that can similarly be transduced into a cellular response (*1, 2*). Biochemical and mechanical signals do not act in a segregated fashion, but instead can function complementarily (*3*). Both signal types can converge on the same signaling cascades and cellular processes, and impact each other to instigate mechanochemical feedback. Unraveling the interplay between biochemical and mechanical signals, and elucidating their underlying hierarchical relationships, is crucial for a comprehensive understanding of how cells integrate the information they receive.

The transduction of forces into cellular reponses relies on a repertoire of mechanosensor proteins. In epithelia, a prominent mechanosensory role is fulfilled by adherens junctions (AJs) (*4, 5*). These adhesion complexes connect epithelial cells to each other and concomitantly sense tensile forces exerted between cells, either originating from tissue-scale morphogenetic movements or from local cell shape changes. AJs are large multiprotein complexes centered around intercellular trans-dimers of E-cadherin that connect to the actin cytoskeleton through β-catenin and α-catenin, the latter of which undergoes force-dependent conformation changes (*6–8*). As a result, the composition of the AJ and the downstream signaling it instigates depend on the amount of intercellular force (*9*). Targeted approaches identified a number of signaling cascades that are controlled by E-cadherin mediated mechanotransduction, including Hippo and Wnt signaling (*10–12*). Accumulating evidence also links forces to extracellular signal regulated kinase (ERK) signaling, implicating mechanical regulation of ERK activity in key physiological processes, from collective cell migration and tissue branching in morphogenesis to cell division and death regulation in tissue homeostasis (*13–17*). Nevertheless, we incompletely understand how this force-induced signaling intersects with the regulation of these signaling cascades by biochemical factors.

Mechanical forces experienced by epithelial tissues can be modeled by application of mechanical stretch to epithelial cultures. Here, we performed unbiased analyses of strain-induced signaling in epithelial monolayers using phosphoproteomics and RNA sequencing, which identified ERK activation by the upstream epidermal growth factor (EGF) receptor (EGFR) as most prominently induced pathway. Because EGFR-ERK signaling is well-known to be instructed by EGF-family ligands, we utilized this finding to unravel the mechanistic nature of mechanochemical crosstalk in epithelial cells. We found that mechanical strain is transduced by AJs to enhance EGFR activity, which is in line with reports that link forces on E-cadherin to the activation of EGFR (*15, 18–20*). Whereas forces on E-cadherin have been reported to act on EGFR itself (*20*), here we demonstrate that intercellular forces transduced by E-cadherin activate EGFR by inducing the shedding of its ligands. As such, our data reveal how forces can modulate external biochemical signals, and exemplify that biochemical and mechanical signals are not necessarily separate inputs that converge on cellular signaling pathways, but can function conjuctively in one single linear signaling cascade.

## Results

### ERK signaling dominates the mechanical strain induced phosphorylation landscape in epithelia

To investigate how mechanical strain impacts intracellular signaling pathways in epithelia, we applied equibiaxial stretch to epithelial cell monolayers and subjected these to phosphoproteomic and transcriptomic analyses. Hereto, we adapted a previously established pneumatically controlled cell stretcher (*21*) to a 6-well format to obtain sufficient protein yield for biochemical analyses (Fig. 1A; fig. S1, A and B). We first mapped strain-induced phosphorylations, because this type of posttranslational modification is both dynamically regulated and most abundantly present (*22, 23*). We labeled proteins in MDCK cells using stable isotope labeling by amino acids in cell culture (SILAC), and subjected monolayers to 18% static substrate stretch for 30 minutes (or left unstretched), thus focusing on immediate responses to strain instead of long-term adaptations. Phosphopeptide ratios were subsequently determined by liquid chromatopgraphy-tandem mass spectrometry (LC-MS/MS) (Fig. 1A), which identified 7795 unique phosphopeptides, 559 of which were differentially present in strained vs. non-strained conditions (313 overrepresented upon strain, 246 underrepresented upon strain) (Fig. 1B; data file S1). Unbiased pathway analysis of the 559 regulated phosphopeptides (data file S2) revealed pathways related to cytoskeleton remodeling and protein synthesis to be altered, in line with mechanical load being known to modulate these processes (*24–26*). In addition, this analysis identified the MAPK signaling pathways to be substantially induced upon mechanical strain, indicating that strain impacts this signal transduction pathway that is classically considered to be induced by biochemical signals. In line with this, the activating tyrosine phosphorylations of ERK1 and ERK2 were amongst the phosphopeptides that showed the greatest increase (Fig. 1B). Furthermore, when we analyzed the regulated phosphopeptides for a consensus within their phosphomotifs, the most enriched motifs within the overrepresented phosphopeptides upon strain are all identified as variants of S/T-P sites, which are ERK kinase substrate motifs (Fig. 1, B to D). In contrast, the phosphopeptides that are underrepresented upon strain contain only few S/T-P sites and are dominated by Serine residues preceded by basic amino acids (Lysine and/or Arginine) (Figs. 1, C and E). Screening all phosphomotifs for the more specific ERK target motif P-x-S/T-P reveals its enrichment among phosphopeptides increased by strain (22.4% (70/313) of increased phosphopeptides vs. 12.6% (982/7795) of all detected phosphopeptides and 5.3% (13/246) of decreased phosphopeptides). As such, these analyses together identify ERK signaling as major pathway activated in epithelial monolayers subjected to mechanical strain.

**Fig. 1.**
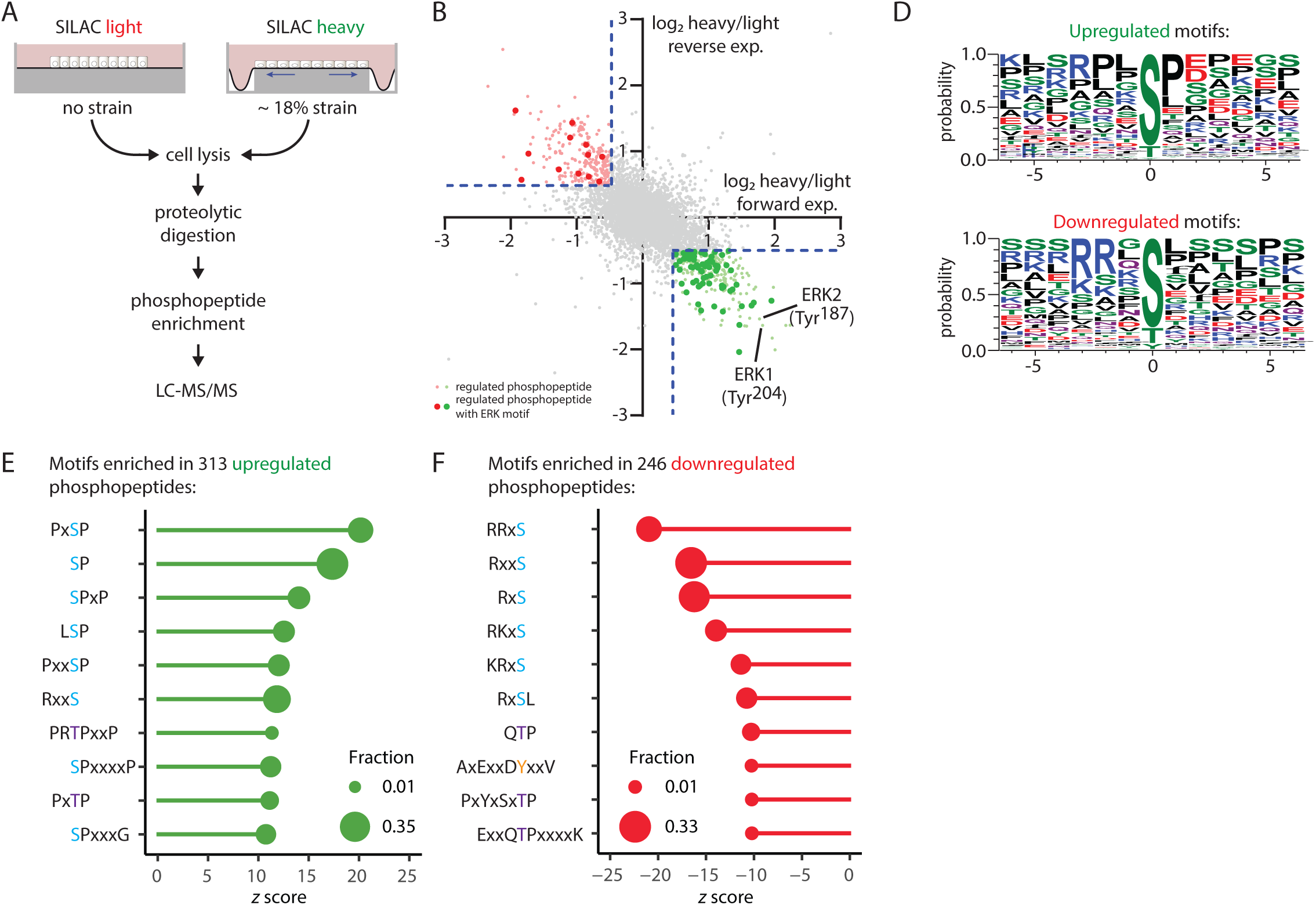
Identification of phosphorylation events regulated by mechanical strain. **(A)** Workflow for identifying mechanical strainregulated phosphorylation events. Differentially SILAC-labeled MDCK cells were subjected to 18% mechanical strain for 30 minutes or left unstrained Cells were lysed and phosphopeptides were isolated and quantified by LC-MS/MS. The forward experiment is depicted and the reverse experiment with SILAC labels swapped was also performed to control for labeling effects. **(B)** Phosphopeptide ratios in forward (x-axis) and reverse experiments (y-axis). Each datapoint represents one phosphopeptide. Green phosphopeptides were overrepresented upon mechanical strain; red phosphopeptides underrepresented. Bold datapoints indicate phosphopeptides containing an ERK target motif (P-x-S/T-P). The activating tyrosine phosphorylations of canine ERK1 (Tyr^204^) and ERK2 (Tyr^187^) are indicated. **(C)** Amino acid probabilities within phosphopeptides that increased or decreased upon the application of mechanical strain. **(D and E)** Phosphomotif enrichment analysis of phosphopeptides that increased (D) or decreased (E) upon strain, ranked by *z* score of each motif and indicating the number of times corresponding motifs were identified as a fraction of the total number of increased or decreased phosphopeptides. Phosphoproteomics data represent n = 1 independent experiment (1 forward experiment, 1 reverse experiment).

### Mechanical strain induces an ERK-mediated transcriptional response

To address whether strain-induced signaling pathways propagate to long-term cellular responses in epithelial cells and potentially validate the role of ERK signaling herein, we investigated the transcriptional response to mechanical strain. MDCK monolayers were subjected to static stretch or left unstretched for 2 hours and transcriptional activity was assessed by RNA sequencing. A total of 17,263 mRNAs was identified by sequencing, 267 of which were differentially present in strained vs. non-strained conditions (192 elevated upon strain, 75 reduced) (Fig. 2A, data file S3). We next performed unbiased analyses to identify upstream regulatory elements of strain-regulated transcripts, focusing on enrichment of either general regulatory signatures (Fig. 2B) or specific transcription regulator signatures (Fig. 2C). The former predominantly identified signaling induced by growth factors or cytokines, with EGF-induced signaling being the most significantly enriched signature regulated by strain (Fig. 2B). Concentrating the analysis of upstream regulators on transcription (co-)factors identified several transcription regulators that are well known to be regulated by mechanical signals, including YAP/TEAD and SRF/MRTFA/MRTFB (Fig. 2C). Besides these expected transcriptional responses, the top regulated transcription factors mainly constitute proteins involved in cellular stress response (TP53, NUPR1, JUN, HIF1A) and proteins known to be controlled by ERK signaling (CREB1, MYC, FOXL2, SP1, EGR1, ELK1). To corroborate this preeminence of strain-regulated ERK-controlled signatures further, we specifically analyzed the transcriptional response of 249 ERK targets, based on published transcriptional analysis of MAPK pathway activity (Fig. 2A, data file S4) (*27–29*). These ERK targets were overrepresented in increased transcripts, constituting 10.4% (20 of 192) of the transcripts within this group, as compared to 1.4% (229 of 16.996) within the non-regulated transcripts. As such, these findings demonstrate that mechanical strain results in the induction of transcription of ERK targets, and concur with the phosphoproteomic analysis that shows strain activates ERK signaling in epithelial monolayers.

**Fig. 2.**
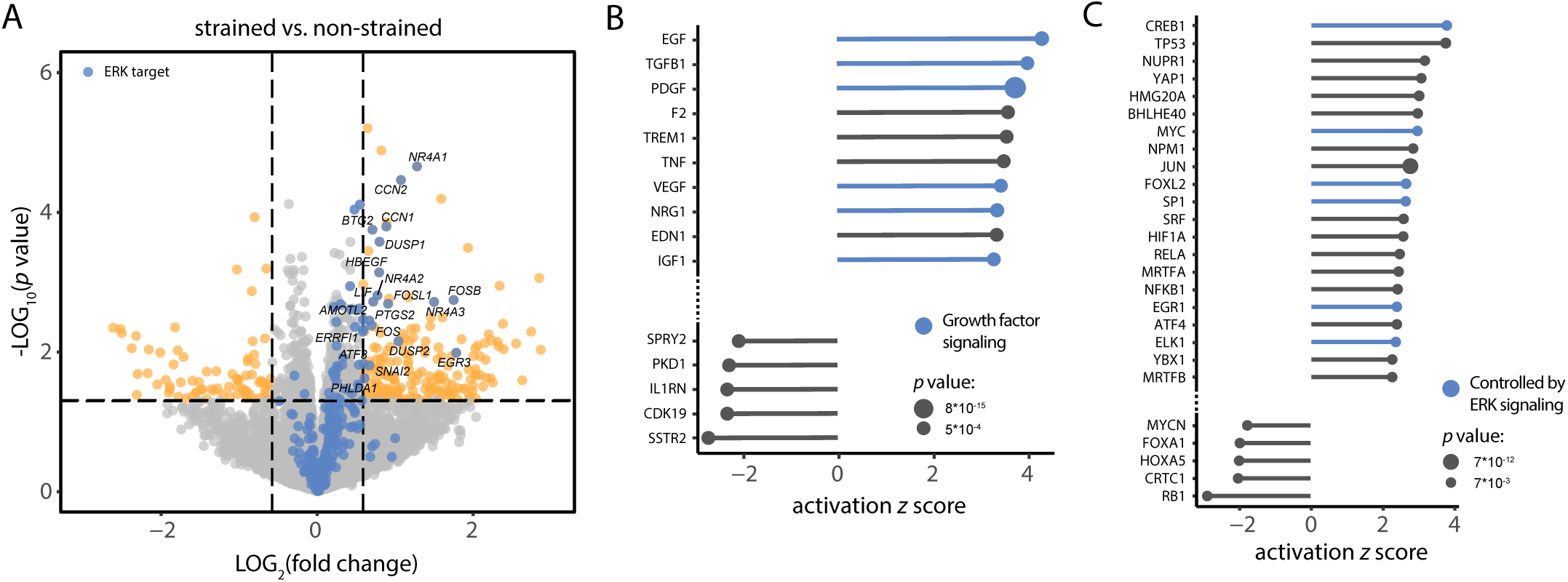
Transcriptional responses to mechanical strain. **(A)** Volcanoplot of transcripts identified in MDCK cells that were either subjected to 18% mechanical strain (2 hours) or left unstrained, showing *p* value (of technical quadruplicates) as a function of the fold change in the strained condition. Transcripts differentially expressed upon mechanical strain are shown in orange; transcripts known to be regulated by ERK in blue (see also data file S4). **(B)** Identification of regulatory pathways upstream of mechanical strain-regulated transcripts using Ingenuity Pathway Analysis. Shown are pathways with the 10 highest and 5 lowest activation *z* scores. Regulatory pathways controlled by growth factors are shown in blue. **(C)** Prediction of upstream transcriptional regulators of mechanical strain regulated transcripts using Ingenuity Pathway Analysis. Shown are transcriptional regulators with the 21 highest and 5 lowest activation *z* scores. Transcriptional regulators controlled by ERK signaling are shown in blue.

### E-cadherin mechanotransduction induces EGFR-ERK signaling at the receptor level

Our forgoing untargeted analyses identified ERK signaling as prominent signaling hub activated in epithelial cells by strain. Validation experiments showed that the strain-induced increase of phosphorylated ERK1 and ERK2 (pThr^202^/pTyr^204^) (pERK) levels can be readily detected by Western blot (Fig. 3, A and B,). The total amount of ERK was unaffected by strain, making tubulin loading controls a suitable proxy for normalization to total protein (Fig. 3B, quantified in fig. S2A). Screening additional cell lines revealed that the strain-induced increase in pERK consistently occurred across human epithelial cell lines (fig. S3). The pERK induction was considerably potentiated in serum-starved conditions that reduced basal levels of pERK (fig. S4, A to C), and these conditions were therefore employed in further experiments. Next, we ventured to assess how mechanical strain impinges on ERK signaling. One major means of ERK activation occurs downstream of EGFR (Fig. 3A), which was previously described to be impacted by various types of mechanical stimuli (*13, 16*). Western blot analysis showed that the activating trans-autophosphorylation of Tyr^1068^ on EGFR is induced upon straining MDCK cells (Fig. 3C), as well as the activating phosphorylations of MEK1 and MEK2 (Ser^217^ and Ser^221^ MEK1/2) that connect EGFR activity to ERK phosphorylation (Fig. 3B). Similar to the effect of strain on ERK, the induction of EGFR and MEK1/2 was observed specifically in their phosphorylated forms with no change in total protein amounts (fig S2, B to C). To directly assess the contribution of EGFR to strain-induced ERK activation, we first inhibited EGFR with the pan-human EGFR (HER) inhibitor Afatinib, which blocks EGFR kinase activity. Afatinib pretreatment reduced the amount of pERK in non-strained cells and largely prevented it to be increased by mechanical strain (Fig. 3D). Some strain-induced pERK increase remained in Afatinib-treated cells, which is likely due to EGFR-independent activation of ERK, because no residual strain-induced EGFR activation was observed in Afatinib-treated cells (Fig. 3E). To corroborate this result further, we used a different class of EGFR inhibitors, which do not block EGFR kinase activity directly, but rather compete with ligand binding to the EGFR extracellular domain. Because Cetuximab and Panitumumab proved to be ineffective in blocking EGF-induced ERK activation in canine MDCK cells (fig. S5), we opted to perform this experiment using human epithelial DLD1 cells, which also displayed ERK regulation by mechanical strain (fig. S3). Pretreatment of DLD1 cells with either EGFR blocking antibody strongly prevented strain-induced activation of ERK (Fig. 3F). Taken together, these data demonstrate that mechanical strain-induced ERK signaling in epithelial cells relies on both EGFR kinase activity as well as EGFR ligand binding.

**Fig. 3.**
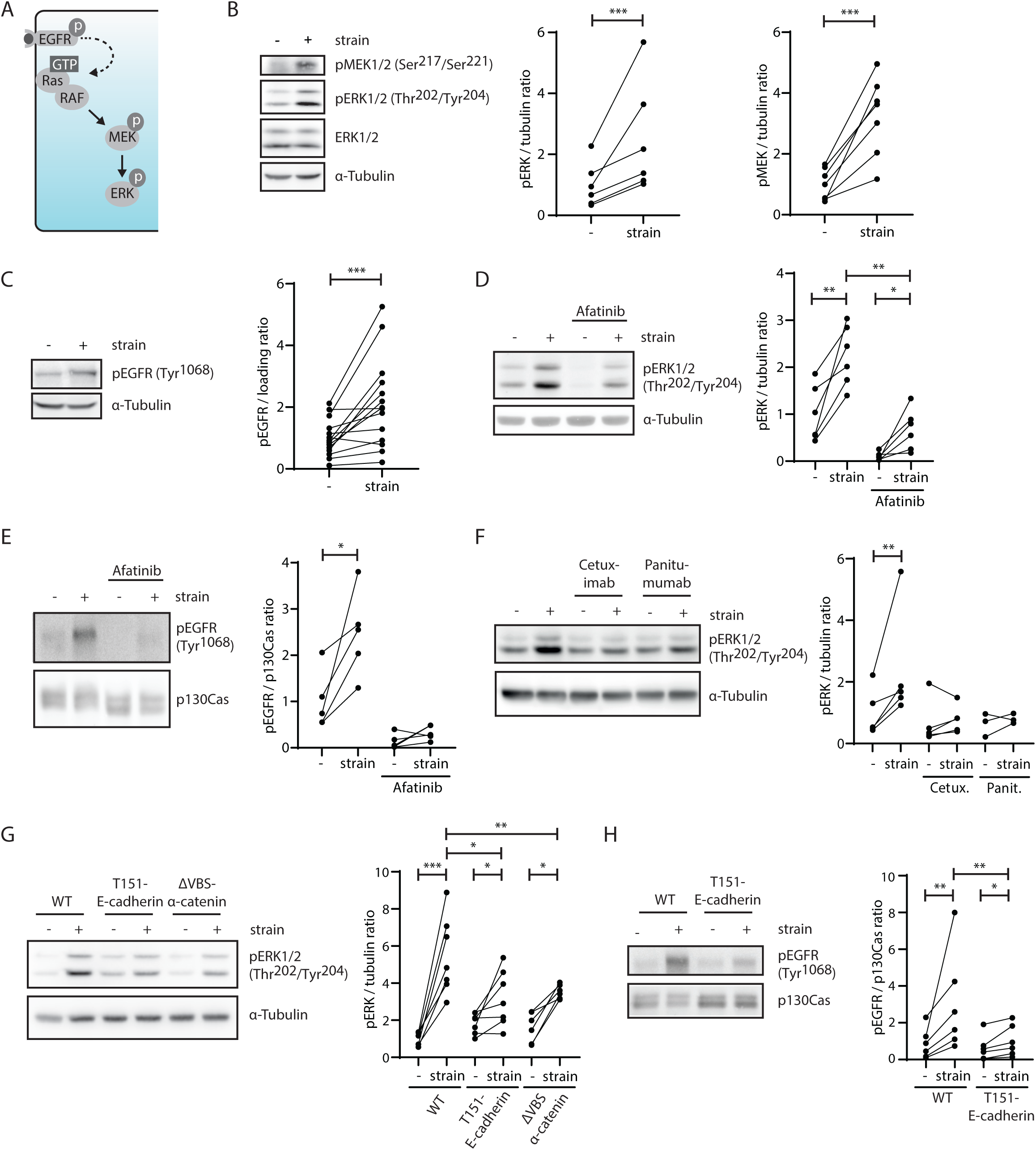
AJ mechanotransduction activates EGFR-ERK signaling at the level of EGFR. **(A)** Schematic representation of the EGFR-ERK signaling pathway. Ligand binding to the EGFR extracellular domain induces the sequential activation of EGFR, Ras, RAF, MEK and ERK. **(B and C)** Western blot of lysates from non-strained and strained (15 minutes) MDCK cells, probed for MEK1 and MEK2 phosphorylated on Ser^217^ and Ser^221^ [pMEK1/2 (Ser^217^/Ser^221^)], ERK1 and ERK2 phosphorylated on Thr^202^ and Tyr^204^ [pERK1/2 (Thr^202^/Tyr^204^)], total ERK1/2 and α-Tubulin. Quantifications show ratios of pERK/α-Tubulin (n = 6 independent experiments), pMEK/α-Tubulin (n = 7 independent experiments) and pEGFR/loading control (either p130Cas or α-Tubulin) (n = 13 independent experiments), normalized to the mean ratio of the non-strained control. Normalization of pERK to total ERK and pMEK to total MEK shows comparable induction of pERK and pMEK by strain, respectively (see fig. S2Aand S2B). **(D and E)** Western blot of lysates from non-strained and strained (15 minutes) MDCK cells, in the presence or absence of Afatinib (1 μM), and probed for pERK1/2 (Thr^202^/Tyr^204^), α-Tubulin, pEGFR (Tyr^1068^) and p130Cas as indicated.Graphs show the ratio of pERK/α-Tubulin (n = 6 independent experiments) and pEGFR/p130Cas (n = 5 independent experiments), normalized to the mean ratio of the non-strained, non-treated control. **(F)** Western blot of lysates from non-strained and strained (15 minutes) DLD1 cells, in the presence or absence of Cetuximab (5 μg/mL) or Panitumumab (20 μg/mL), and probed for pERK1/2 (Thr^202^/Tyr^204^) and α-Tubulin. The quantification shows the ratio of pERK/α-Tubulin, normalized to the mean ratio of the non-strained, non-treated control (n = 5 independent experiments). **(G)** Western blot of lysates from MDCK, MDCK-T151-E-cadherin and MDCK-ΔVBS-α-catenin cells, either strained (15 minutes) or unstrained, and probed for pERK1/2 (Thr^202^/Tyr^204^) and α-Tubulin. The quantification shows the ratio of pERK/α-Tubulin in WT MDCK (n = 7 independent experiments), MDCK-T151-E-cadherin (n = 7 independent experiments) and MDCK-ΔVBS-α-catenin cells (n = 6 independent experiments), normalized to the mean ratio of the non-strained, WT MDCK control. **(H)** Western blot of lysates from MDCK and MDCK-T151-E-cadherin cells, either strained (15 minutes) or unstrained, and probed for pEGFR (Tyr^1068^) and p130Cas. The quantification shows the ratio of pEGFR/p130Cas, normalized to the mean ratio of the non-strained, WT MDCK control (n = 6 independent experiments). All statistical tests shown are paired ratiometric t-tests. * p < 0.05, ** p < 0.01, *** p < 0.005.

In epithelia, AJs fulfill an important function in the sensing and transduction of intercellular forces (*4, 5*). Because AJ mechanotransduction is implicated in regulation of EGFR-ERK (*18–20*), we tested whether it is directly responsible for strain-induced activation of EGFR-ERK in MDCK epithelial cells. We used two different genetic approaches to attenuate AJ mechanotransduction. First, we used MDCK cells with inducible expression of truncated E-cadherin that lacks its extracellular domain (T151-E-cadherin), which replaces endogenous E-cadherin upon induction of its expression (*30*). T151-E-cadherin cannot transmit forces between cells, and consequently mechanical stretch cannot exert direct tensile forces on E-cadherin (*31*). Second, we utilized MDCK cells in which endogenous α-catenin was replaced by α-catenin in which the region that becomes exposed upon forces and consequently can recruit additional proteins, including Vinculin, is mutated (ΔVBS-α-catenin). Cells expressing this mutant maintain their AJ composition in the low-force state even when intercellular force is exerted on them (*32*). Both these mutant cell lines displayed slightly elevated ERK activity in non-strained conditions and significantly decreased ERK activity in strained conditions as compared to wild type MDCK cells (Fig. 3G), indicating that mechanical strain is transduced by AJs to activate ERK signaling.

Altogether, these data demonstrate that mechanical strain activates EGFR-ERK signaling at the receptor level and that AJ mechanotransduction plays an important role herein. Intriguingly, although residual strain-induced ERK activation remains in the T151-E-cadherin mutant, the increase in phospho-EGFR after strain was almost fully abrogated in this mutant cell line (Fig. 3H). As such, despite AJ mechanotransduction being essential for strain-induced EGFR activation, additional mechanosensitive mechanisms may contribute to activation of ERK independently of E-cadherin and EGFR.

### Mechanical strain activates EGFR-ERK by inducing ADAM-mediated ligand shedding

In search of a mechanistic link between EGFR and force transduction by the AJ complex, we performed proximity labeling of the AJ complex to identify a potential association of proteins linked to the regulation of EGFR activity. We fused APEX2 to the cytosolic C-terminus of E-cadherin (Fig. 4, A and B), therewith enabling H_2_O_2_-induced biotinylation of proteins in proximity of E-cadherin that can be readily identified by LC-MS/MS (Fig. 4, B and C). Comparison of E-cadherin-APEX2 to an APEX2-NLS control showed enriched biotinylation of the core AJ proteins α-catenin, β-catenin, p120-catenin and E-cadherin itself (Fig. 4C, data file S5). Apart from these core AJ proteins, one of the proteins identified to be in close proximity to E-cadherin was the metalloproteinase ADAM17 (also known as TACE, tumor necrosis factor-converting enzyme), which catalyzes the shedding of EGFR ligands (*33*). Because our experiments show that EGFR-ERK activation by strain requires ligand binding to EGFR (Fig. 3F) but occurs in the absence of exogenously supplied ligand (fig. S4, A to C), we hypothesized that E-cadherin mechanotransduction may impinge on ADAM17 to induce EGFR ligand shedding and concomitant EGFR activation. To test this hypothesis, we assessed the contribution of ADAM17 and related ADAM proteins to stretch-induced EGFR-ERK activation using the ADAM inhibitor TAPI-2. Pretreatment of cells with TAPI-2 largely prevented the increase in pERK levels by mechanical strain (Fig. 4D). The slight remaining ERK activation is likely caused by the activity of an EGFR-independent input into ERK activation, because EGFR activity was abolished upon TAPI-2 pretreatment in both non-strained and strained cells (Fig. 4E). Inhibition of ADAM activity using the alternative inhibitor Marimastat resulted in a similar loss of strain-induced pERK increase (fig. S6).

**Fig. 4.**
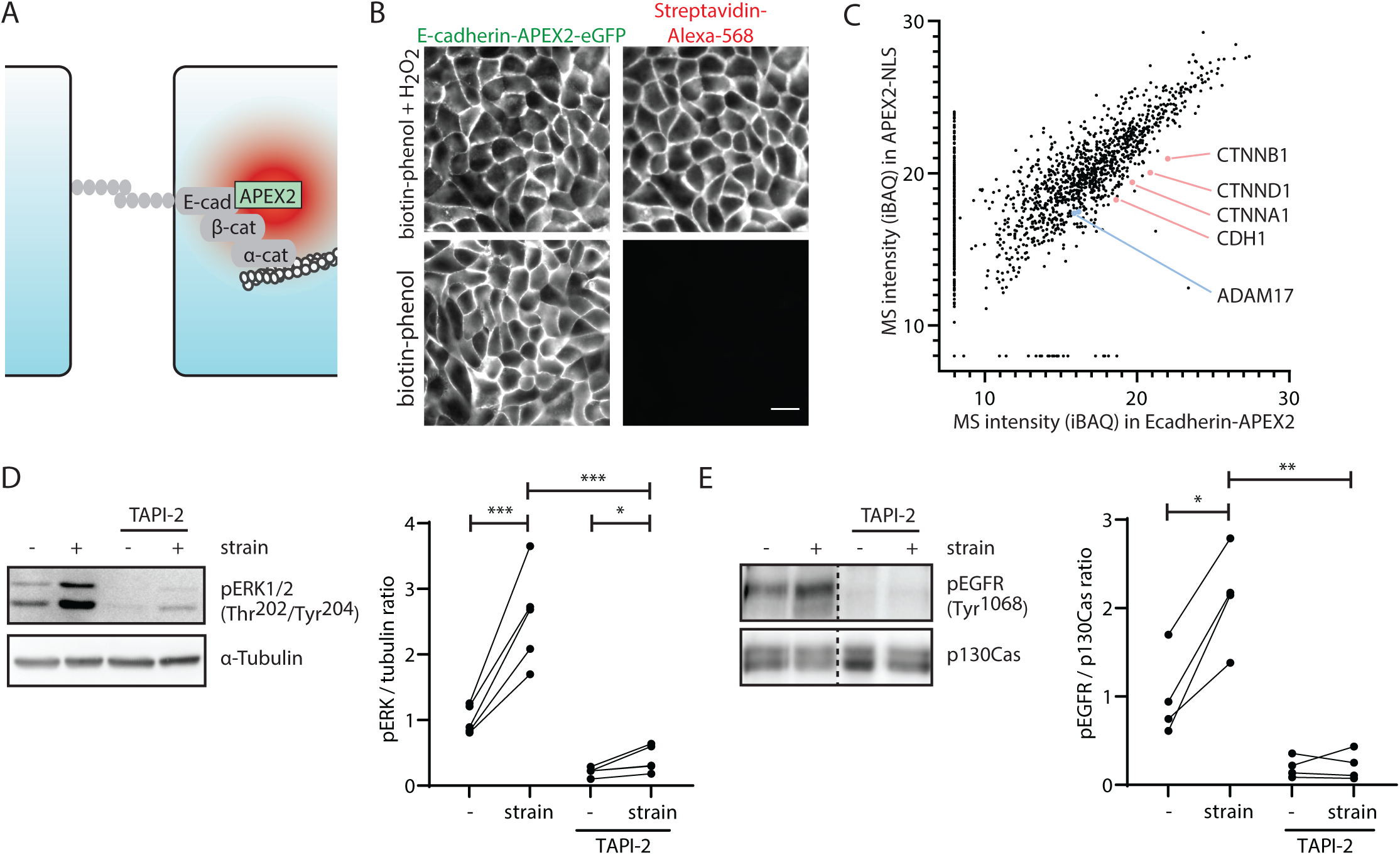
Mechanical strain induces ADAM-dependent EGFR ligand shedding. **(A)** Schematic representation of APEX2-mediated biotinylation of proteins in proximity of E-cadherin. APEX2 fused to the C-terminus of E-cadherin generates a gradient of biotin-phenoxyl (red) that covalently binds proteins. The high reactivity of biotin-phenoxyl ensures labeling only of proteins in close proximity to E-cadherin. **(B)** Visualization of biotinylated proteins in MDCK-E-cadherin-APEX2-GFP cells, following treatment with biotin-phenol and H_2_O_2_ (or biotin-phenol only as negative control) with Alexa Fluor 568-conjugated Streptavidin. Images are a representation of n = 3 independent experiments. Scalebar, 20 μm. **(C)** Scatter plot of iBAQ intensities of proteins identified by mass spectrometry in Streptavidin pull-downs of both MDCK-E-cadherin-APEX2-GFP (x-axis) and MDCK-mNeon-APEX2-NLS control cells (y-axis). Proteins that were only identified in one sample are shown with an imputed iBAQ intensity of 8 in the negative sample. Indicated are the core AJ proteins E-cadherin (CDH1), β-catenin (CTNNB1), α-catenin (CTNNA1) and p120-catenin (CTNND1), as well as ADAM17. N = 1 independent experiment. **(D and E)** Western blot of lysates from non-strained and strained (15 minutes) MDCK cells, in the presence or absence of TAPI-2 (50 μM), and probed for phosphorylated ERK [pERK (Thr^202^/Tyr^204^)], α-Tubulin, pEGFR (Tyr^1068^) and p130Cas as indicated. Graphs show the ratio of pERK/α-Tubulin (n = 5 independent experiments) and pEGFR/p130Cas (n = 4 independent experiments), normalized to the mean ratio of the non-strained, non-treated control. * p < 0.05, ** p < 0.01, *** p < 0.005.

Our data show that ADAM enzymatic activity is required for strain-induced EGFR-ERK activation and suggest an instructive function of ADAM enzymes in this process, with ADAM function being regulated by intercellular forces transduced by E-cadherin. Alternatively, these data may be explained by a merely permissive function of ADAM proteins, in which continuous ADAM activity provides a non-rate limiting supply of EGFR ligand and E-cadherin mechanotransduction acts by another mechanism to activate EGFR-ERK. To discriminate between these possibilities, we directly analyzed the effect of mechanical strain on ADAM activity. Therefore, we generated an ADAM activity probe, consisting of the full-length EGFR ligand EREG tagged with an mNeon fluorophore at its cytosolic C-terminus and an mScarlet fluorophore at its extracellular N-terminus (Fig. 5A). Changes in ADAM activity will elicit altered mScarlet/mNeon ratios at the plasma membrane, because the mScarlet moiety is being shed along with the EREG ectodomain. In control conditions, both mNeon and mScarlet were readily observed at the plasma membrane (Fig. 5, B and C). Upon mechanical strain, mScarlet intensity at the plasma membrane was reduced, whereas mNeon intensity remained unaltered (Fig. 5, B and C). Quantification of the mScarlet/mNeon ratio showed that the presence of the uncleaved probe at individual cell-cell contacts was significantly decreased after mechanical strain (Fig. 5 C and D, fig. S7A). These findings indicate that ADAM activity is in itself increased when cells experience mechanical strain. Strain-induced cleavage of the mScarlet-EREG-mNeon probe was inhibited by pretreatment of cells with TAPI-2, validating the specific role of ADAM activity in this cleavage (Fig. 5E). Furthemore, a control probe consisting of mScarlet and mNeon fused to the unrelated protein NECL5, which is shed in a thrombin-dependent manner (*34*), was not affected by mechanical strain (fig. S7 B and C). Finally, cells in which endogenous E-cadherin is replaced with T151-E-cadherin do not show significant strain-induced cleavage of mScarlet-EREG-mNeon, confirming that AJs are essential for relaying mechanical strain to control ADAM activity (Fig. 5F).

**Fig. 5.**
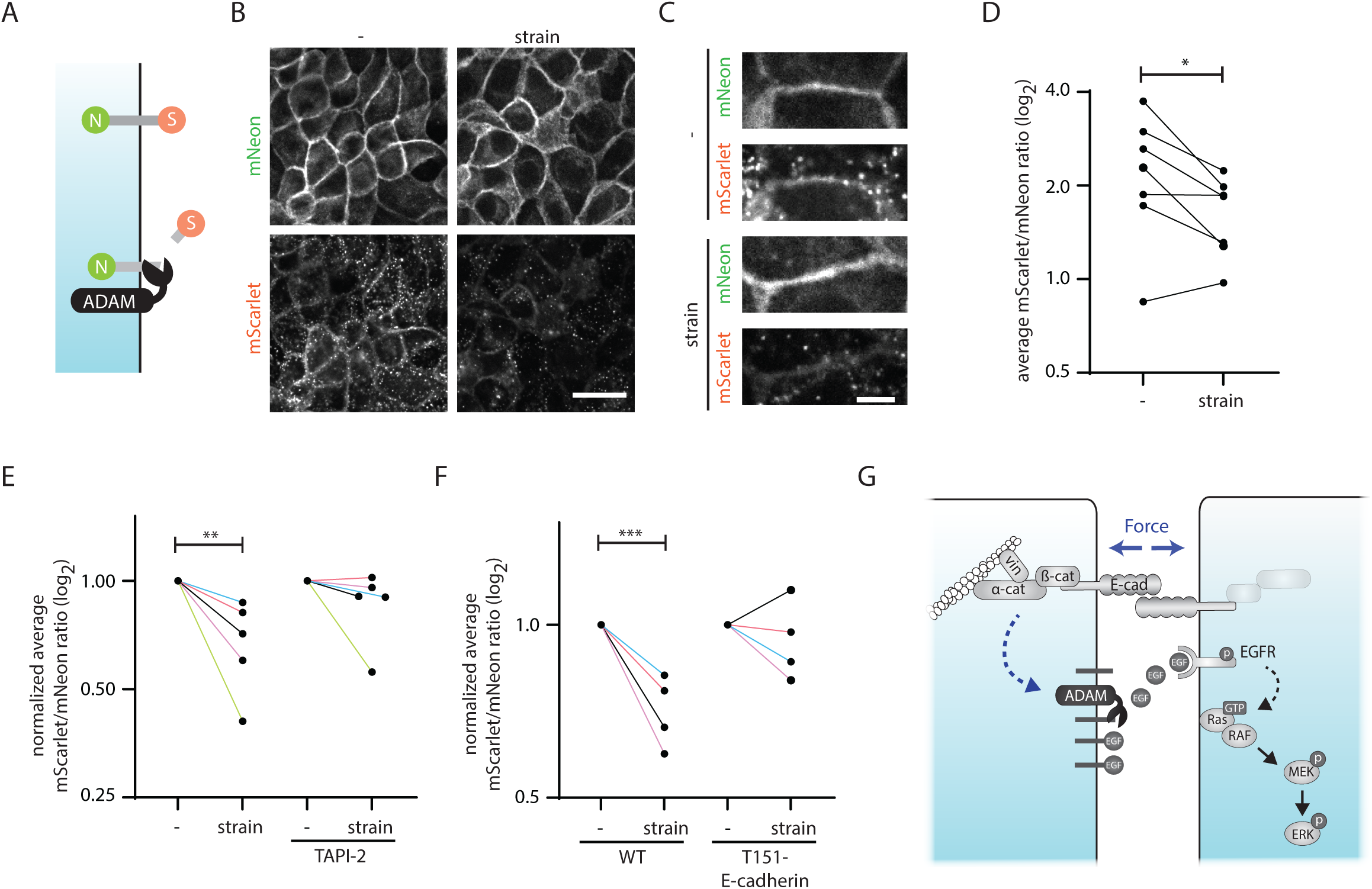
E-cadherin mechanotransduction instructs ADAM activity. **(A)** Schematic representation of the ADAM activity probe. mNeon (green) and mScarlet (orange) fluorescent proteins were fused to the cytosolic C-terminus and extracellular N-terminus of EREG, respectively. ADAM proteins cleave EREG in the extracellular domain, therewith shedding the mScarlet. **(B)** Representative example of fixed MDCK cells expressing the ADAM activity probe, either non-strained or subjected to mechanical strain (15 minutes). Images are a representation of = 7 independent experiments. Scalebar, 20 μm. **(C)** High magnification views of individual cell-cell contacts. Scalebar, 5 μm. **(D)** Quantification of junctional mScarlet/mNeon ratios of MDCK cells expressing the ADAM activity probe, either non-strained or subjected to mechanical strain (15 minutes) (n = 7 independent experiments). Each data point represents the mean ratio of an independent experiment, in which ≥ 65 cell-cell junctions were quantified across at least two field of views per condition. Statistical tests are paired ratiometric t-tests. **(E)** Quantification of the effect of mechanical strain (15 minutes) on junctional mScarlet/mNeon ratios of MDCK cells expressing the ADAM activity probe, in the presence or absence of TAPI-2 (n = 5 independent experiments). Each data point represents the mean ratio of an independent experiment, normalized to non-strained control. ≥ 85 cell-cell junctions across at least three fields of view were quantified per condition in each individual experiment. Statistical tests are unpaired t-tests. **(F)** Quantification of the effect of mechanical strain (15 minutes) on junctional mScarlet/mNeon ratios of MDCK cells (WT) or MDCK-T151-E-cadherin cells expressing the ADAM activity probe (n = 4 independent experiments). Each data point represents the mean ratio of an independent experiment, normalized to non-strained control. ≥ 160 cell-cell junctions across five fields of view were quantified per condition in each individual experiment. Statistical tests are unpaired t-tests. **(G)** Schematic model of the mechanism by which mechanical strain activates EGFR-ERK signaling in epithelia. Our findings demonstrated that E-cadherin-based AJs transduce strain to induce ADAM-mediated cleavage of EGFR ligands, allowing them to bind to EGFR family receptors and instigating EGFR-ERK signaling. This may act in concert with other mechanisms relaying forces to ERK, see *Discussion*. Note that cleaved EGFR ligands may not only act in a paracrine manner (as shown) but also autocrine manner. * p < 0.05, ** p < 0.01, *** p < 0.005.

Taken together, our data indicate that ADAM activity is modulated by intercellular forces acting on the AJ, thereby inducing shedding of EGFR ligands and concomitant EGFR-ERK activation.

## Discussion

To regulate their behavior, cells integrate biochemical and mechanical information that they receive, yet the interplay between these types of signals remains incompletely understood. Here, we employed phosphoproteomic and transcriptomic analyses of MDCK cells exposed to stretch to uncover mechanically controlled signal transduction in epithelia and identify potential hubs of mechanochemical crosstalk. This unveiled that forces induced by mechanical stretch are transduced by E-cadherin adhesions to instruct ADAM-mediated shedding of EGFR ligands, consequently activating EGFR-ERK signaling (Fig 5G). Our findings therewith introduce a novel paradigm in the interplay between intercellular force and receptor ligand-induced signal transduction. Previously, the transduction of intercellular forces was shown to impinge on intracellular signaling cascades either in a manner independent of biochemical ligands or by altering ligand sensitivity (*2*). Our results now demonstrate transduction of forces directly modulates the presence of the biochemical signal to regulate downstream pathway activity, thereby illustrating how intercellular force and biochemical ligands can function within a single linear cascade.

Our unbiased analyses of stretched epithelial cells most prominently shows the induction of EGFR-ERK signaling, with a substantial amount of regulated phosphomotifs and transcripts linked to activity of this pathway. This implies EGFR-ERK signaling to be either the prominent mechanoresponsive pathway in epithelial cells or the mechanoresponsive pathway that propagates into the most pervasive set of cellular changes. The prevalence of EGFR-ERK activation following elevation of intercellular forces may further depend on cellular and mechanical conditions, including cell type and density, as well as force magnitude and duration. In line with this, although EGFR-ERK is broadly observed to be induced by mechanical signals (*13, 16*), the extent of this response varies across cell types and mechanical stimuli (*35–40*).

ERK signaling is well established to be induced by a variety of mechanical stimuli, in which a myriad of mechanosensing mechanisms are implicated (*13, 16*). It becomes evident from our data that the transduction of mechanical strain to ERK signaling in epithelial monolayers is established by E-cadherin adhesions via regulation of EGFR. Because inhibition of E-cadherin mechanotransduction strongly attenuates but not fully prevents strain-induced phosphorylation of ERK, additional mechanosensitive mechanisms may contribute to achieving full-blown activation of ERK. For instance, outside-in signaling by integrins is implicated in the mechanical activation of ERK independently of EGFR signaling (*41*), and mechanosensitive influx of calcium activates ERK through noncanonical EGFR endocytosis and signaling (*14, 42*). Future research may elucidate the interplay between these mechanosensitive mechanisms, and reveal how they may relay different types of mechanical information to the regulation of ERK. Notably, our gene expression analysis indicates that strain induces transcriptional changes linked to upstream pathways beyond EGFR-ERK signaling. These changes include both previously unrecognized transcriptional responses to mechanical signals as well as previously characterized force-regulated changes (for example, targets of YAP/TEAD and SRF-MRTF), which in part are also linked to AJ-dependent mechanotransduction (*43, 44*).

By visualizing and manipulating ADAM activity, we provide evidence that force transduction by E-cadherin adhesions controls EGFR activity by regulating ligand presence mediated by this sheddase. It has previously been established that E-cadherin also physically interacts with EGFR, and dissociation of this complex is needed for EGFR dimerization and subsequent trans-autophosphorylation essential for its activation (*20, 45*). It was reported that dissociation of the E-cadherin/EGFR complex is induced by ligand binding, as well as by external application of mechanical stretch (*20*). Mechanical regulation of ADAM-mediated ligand shedding and disassembly of the E-cadherin/EGFR complex could thus act synergistically to stimulate EGFR-ERK pathway activation. Alternatively, these mechanisms may function interdependently, given that E-cadherin-EGFR complex disassembly occurs upon receptor binding to ligands, and ADAM-mediated shedding could therefore contribute to ligand availability and complex disassembly.

Understanding how mechanosensing by E-cadherin is coupled to the regulation of ADAM sheddases warrants further investigation. Although proximity labeling specifically identified ADAM17 to be in proximity of E-cadherin, other ADAM sheddases that may be inhibited by TAPI-2 may similarly contribute to stretch-induced ligand shedding. We could not biochemically demonstrate an interaction between E-cadherin and ADAM17 proteins, suggesting E-cadherin controls ADAM through intermediary pathways rather than a direct physical interaction. In muscle cells, mechanical strain has been reported to induce ADAM17 activity through Src-mediated phosphorylation (*46–48*), and elevated Src activity at cell-cell contacts has been observed following strain (*49*). In the *Drosophila* gut, a reduction of E-cadherin in apoptotic cells was shown to increase their EGF shedding through a transcriptional induction of its membrane-bound protease (*50*). However, transcriptional regulation is unlikely to explain the observed induction of ADAM, EGFR and ERK activity that follows the application of stretch within minutes. In keratinocytes, weakening of AJs by means of α-catenin depletion leads to enhanced ADAM17-mediated shedding of EGFR ligands (*51*), suggesting that E-cadherin adhesions inhibit ADAM activity and that this may potentially be relieved by intercellular forces.

ERK signaling is implicated in a wide range of force-regulated cellular processes underlying tissue morphogenesis and homeostasis (reviewed in (*13*)). We previously demonstrated that the coordination of collective epithelial cell migration relies on the propagation of ERK activity waves, which is directed by forces transmitted between neighboring cells (*15, 52*). This propagation of ERK signaling was shown to be disrupted upon depletion or inhibition of α-catenin, ADAM17 or EGFR (*15*), suggesting either a permissive or instructive role of both ADAM17 and AJs. We now provide direct evidence that ADAM-mediated EGFR ligand shedding is actively regulated by forces transduced by AJs (Fig. 5). This further implicates this mechanosensitive pathway in the mechanical regulation of collective migration and provides an understanding of the underlying molecular mechanism. Future studies may reveal whether other force-dependent cellular processes also rely on upstream AJ-mediated mechanical activation of ADAM-EGFR-ERK signaling.

The identified linear relationship between mechanical and biochemical signals, wherein forces modulate the availability of receptor ligands rather than altering sensitivity to these signals, has substantial implications for our understanding of tissue regulation by forces. With E-cadherin mechanotransduction affecting ligand shedding, cells are able to autonomously control ERK signaling in response to force, without relying on ligand production by external sources. This cell-autonomous response to forces could, for instance, establish direct control of cell proliferation by changes in forces in an epithelial layer. As such, intercellular forces that rise due to a reduction in cell number within the epithelium (*53*) could directly induce ligand-mediated EGFR activation without requiring the presence of additional external signals. Additionally, E-cadherin mechanotransduction acting on ligand shedding may expand the range of cells capable of responding to intercellular forces. Individual cell populations within a tissue may experience different amounts of forces and exhibit varying degrees of force sensitivity. In this case, cells irresponsive to forces could be supplied with EGFR ligands by neighboring mechanoresponsive cells, which would not be possible if intercellular forces were only to affect EGFR sensitivity to ligands. Given these implications, a linear relationship between mechanical signals and biochemical ligand production clearly enriches cellular control of tissue homeostasis by either signal type.

## Methods

### Plasmids

The lentiviral expression vector containing APEX2 fused to E-cadherin (pLV-CMV-E-cadherin-APEX2-eGFP-IRES-puro) was generated by isolating APEX2 DNA from pCDNA3-APEX2-NES (Addgene, 49386) and introducing this inbetween E-cadherin and eGFP in the pLV-CMV-E-cadherin-eGFP-IRES-puro vector (Johan de Rooij, UMC Utrecht) by InFusion cloning. A 12 amino acid flexible linker (GGGSGGGSGGGS) was introduced inbetween E-cadherin and APEX2 for optimal APEX2 activity. To generate mNeonGreen-APEX2-NLS-IRES-puro, APEX2 was isolated from pCDNA3-APEX2-NES with an NLS integrated in the 3’ PCR primer and together with mNeonGreen DNA isolated from pLV-CMV-H2B-mNeonGreen-IRES-blast (Bas Ponsioen, UMC Utrecht) introduced into pLV-CMV-IRES-puro by InFusion cloning. To generate pCSIIpuro-EREG-ScNeo (Addgene, 209896), which was used to create MDCK-mScarlet-EREG-mNeon, cDNAs of the HA-tag (YPYDVPDYA), mScarlet (Addgene, 85042) and a 8 amino acid flexible linker (GGGSGGGS) were fused in this order and inserted at the 5’ end of the EGF domain of EREG (NCBI CCDS database no. CCDS3564.1) synthesized by GeneArt (Thermo Fisher Scientific). The cDNA of mNeonGreen (*54*) was also synthesized with codon optimization and inserted into the 3’ end of EREG. The cDNA encoding the fusion protein was introduced into the pCSII vector (*55*) by InFusion cloning. To generate pCSIIblast-EREG-ScNeo, which was used to create MDCK-T151-E-cadherin-mScarlet-EREG-mNeon, the pCSIIpuro-EREG-ScNeo was linearized excluding the puromycin cassette by PCR and recircularized with a blasticidin cassette using InFusion cloning. To generate pCSIIpuro-Necl5-ScNeo (Addgene, 209901), cDNAs coding the signal peptide of Necl5 (a.a. 1–28), mScarlet, Necl5 (a.a. 29–408), and mNeonGreen were obtained from pPBbsr2-Necl5-ScNeo (Addgene, 170283) and introduced into ther pCSII vector by ligation.

### Cell lines and culture

Madin-Darby Canine Kidney (MDCK) GII cells and derivatives were cultured at 37°C and 5% CO_2_ in low glucose Dulbecco’s Modified Eagle Medium (DMEM) (Sigma, D5523) containing 10% fetal bovine serum, 1 g/L sodium bicarbonate and 1% penicillin/streptomycin. For SILAC experiments, MDCK cells were grown in low glucose DMEM (Athenaes, 0431) supplemented with 10% dialyzed fetal bovine serum (Silantes, 281001200), 1 g/L sodium bicarbonate, 30 mg/L L-Methionine, 105 mg/L -Leucine and either ^14^N ^12^C - lysine (Sigma, L8662) and ^14^N ^12^C -arginine (Sigma, A8094) (SILAC light) or ^15^N ^13^C -lysine (Silantes, 211604102) and ^15^N_4_^13^C_6_-arginine (Silantes, 201604102) (SILAC heavy). MDCK cells stably expressing doxycycline-repressible truncated E-cadherin (MDCK-T151-E-cadherin) are previously published (*56*) and were cultured in the absence of doxycycline for at least two weeks before experiments. MDCK-α-catenin^−/-^ cells stably expressing ΔVBS-α-catenin (MDCK-ΔVBS-α-catenin) were published previously (*57*). MDCK-E-cadherin-APEX2-GFP, MDCK-mScarlet-EREG-mNeon, MDCK-mScarlet-NECL5-mNeon and MDCK-T151-E-cadherin-mScarlet-EREG-mNeon were generated by lentiviral transduction, and monoclonal lines were obtained by single cell FACS sorting, apart from MDCK-mScarlet-NECL5-mNeon and MDCK-T151-E-cadherin-mScarlet-EREG-mNeon, which are polyclonal sorted lines with naturally uniform expression. DLD1 cells were provided by Boudewijn Burgering (UMC Utrecht) and cultured at 37°C and 5% CO_2_ in RPMI-1640 (Sigma, R0883) containing 10% fetal bovine serum and 1% penicillin/streptomycin. Caco-2 cells were provided by Saskia van Mil (UMC Utrecht) and cultured at 37°C and 5% CO_2_ in high glucose DMEM (Sigma, D6429) containing 1% non-essential amino acid supplement, 10% fetal bovine serum and 1% penicillin/streptomycin. MCF10A cells were provided by Patrick Derksen (UMC Utrecht) and cultured at 37°C and 5% CO_2_ in Ham’s F-12 medium (Sigma, N4888) supplemented with 5% horse serum, 2mM Glutamine, 1% penicillin/streptomycin, 20 ng/mL Epidermal Growth Factor (EGF), 100 ng/mL Cholera toxin, 10 μg/mL Insulin and 1 μM Dexamethasone.

### Application of mechanical strain

6-well stretch devices were adapted from designs described by Simmons et al., (*21*) and generated by attaching a stretchable polydimethylsiloxane (PDMS) membrane (0.25 mm thickness, LSR Sheeting) to a custom made polymethylmethacrylate (PMMA) 6-well plate containing circular apertures, using polypropylene double-sided adhesive tape (Adhesives Research, ARseal). The PDMS membrane was subsequently plasma treated (Harrick Plasma, PDC-002-CE) to enable surface coating. The center of each well was coated with 0.05 mg/mL rat tail Collagen Type 1 (Corning) in 0.1% (v/v) acetic acid (Merck), using a stainless steel ring (Ø 17 mm) placed in the center of each well and plates were sterilized by 1-minute exposure to UV light (Eprom Eraser, LR 123A, Leap Electronic co.,). Cells were plated onto the stretch devices in 400 µL drops of cell suspension at 1.7 × 10^5^ cells/cm^2^ (MDCK, Caco-2 and DLD1) or 1 × 10^4^ cells/cm^2^ (MCF10A). To optimize homogeneity in cell density, MDCK cells were plated in low-calcium (50 µM) DMEM (USbio, D9800) (with the exception of SILAC experiments). For all cell types, seeding medium was replaced by regular culture medium 1-2 hours after plating. Cells were cultured overnight, if applicable in serum-starved conditions (0.5% fetal bovine serum), before application of stretch according to Simmons et al. (*21*). In brief, the cell plate was placed on top of a 6-well PMMA baseplate containing 3 flat-bottom wells and three wells with a central pillar surrounded by a pneumatic compartment (Fig. S1A). Equi-biaxial in-plane strains in the cell monolayer were achieved by vacuum application to the pneumatic compartment using Pressure Controller 2 (Red Dog Research). The correlation between amounts of pressure and strain was determined by direct measurement of the strain of marked PDMS membranes (Fig. S1B) and an air pressure of 90 kPa (resulting in 18% substrate strain) was used for all cell strain experiments.

If applicable, inhibitors were added to the cell culture 30 minutes before stretch application at the following concentrations: Cetuximab (Erbitux, Merck BV): 5 μg/mL, Panitumumab (Vectibix, Amgen): 20 μg/mL, Afatinib (Selleckchem, S1011): 1 uM, TAPI-2 (Santa Cruz, sc-205851): 50 μM, Marimastat (Sigma-Aldrich, M2699): 20 μM.

### Phosphoproteomics

MDCK cells were metabolically labeled with ^14^N_2_^12^C_6_-lysine and ^14^N_4_^12^C_6_-arginine (SILAC light) or ^15^N_2_^13^C_6_-lysine and ^15^N_4_^13^C_6_-arginine (SILAC heavy). Heavy label incorporation was confirmed to be at least 95% before experiments. Cells were plated onto stretch plates (light on non-stretched wells and heavy on stretched wells in the forward experiment, and the opposite for the reverse experiment), stretched for 30 minutes as described above and lyzed in buffer containing 8M urea, 1M ammonium bicarbonate (ABC), 10 mM tris(2-carboxyethyl)phosphine hydrochloride (TCEP), 40 mM 2-chloroacetamide (CAA), 1:50 (w/v) EDTA-free protease inhibitor cocktail (Roche, 12273700), 1% (v/v) phosphatase inhibitor cocktail 2 (Sigma, P5726), and 1% (v/v) phosphatase inhibitor cocktail 3 (Sigma, P0044). To obtain sufficient protein (∼20 mg), 15 sequential runs of two 6-well plates were performed. All lysates were pooled, heavy and light SILAC-labelled samples were mixed and diluted 4-fold with 1 M ABC, upon which trypsin (Worthington) was added (20 ng/mg protein). After protein digestion for 18 hours at 37°C, the peptides were loaded on Sep-pak (Waters, 3CCTC18 cartridge). Titanium dioxide (TiO_2_) beads (ZirChrom) were prepared (10:1 beads:protein) by washing them with 5% NH_3_ (in H_2_O) using centrifugation, followed by loading solution (80% acetonitrile (ACN), 6% trifluoroacetic acid (TFA)). The peptides were eluted from Sep-pak with loading solution and combined with the TiO_2_ beads. After a 5 minute incubation at 37°C, the beads were loaded on an in-house produced C8 stage-tip and the flow-through was discarded after centrifugation. After washing three times with wash buffer (80% ACN, 1% TFA) and once with transfer buffer (80% ACN, 0.5% formic acid (FA)), phospho-peptides were eluted with 200 µL 5% NH_3_ in 30% ACN, into 40 µL 20% FA after which the sample volume was reduced in a SpeedVac concentrator to 20 µL. To acidify the solution, 20 µL of 10% FA was added, before loading the peptides on a C18 stage tip. Peptides were immediately fractionated offline using 5% ACN (pH=10), followed by 10% ACN (pH=10), followed by 50% ACN with 0.1% FA. The volumes of the fractions were reduced in a SpeedVac concentrator and samples were acidified with 0.1% FA. The peptides were separated by liquid chromatography on a 30 cm pico-tip column (75 µm ID, New Objective) with 1.9 µm aquapur gold C-18 material (dr. Maisch). The peptide mixtures were eluted using a 140 minute gradient (7% to 40%, ACN 0.1% FA), delivered by an easy-nLC 1000 (Thermo Scientific), and electro-sprayed directly into a Orbitrap Fusion Tribrid Mass Spectrometer (Thermo Scientific). Detection in ion trap was operated in data-dependent mode with a cycle time of 1 second, in which the full scan over the 350-1,500 mass range was performed at a resolution of 240,000. Most intense ions (intensity threshold of 1.5 ×10^4^) were isolated by the quadrupole and then fragmented with a HCD collision energy of 30%. The maximum injection time of the ion trap was set to 50 milliseconds.

Raw data were analyzed with Maxquant software (*58*) using the *canis lupus familiaris* protein database (Uniprot) with oxidation of methionine, protein N-terminal acetylation and phosphorylation of serine, threonine and tyrosine set as variable modification and carbamidomethylation set as a fixed modification. Protein and peptide false discovery rate cut-off was set to 1%. Phosphopeptides were filtered for reverse hits, standard contaminants, ratio counts > 1 and mass error < 1 ppm in Perseus (v1.5.5.3). Normalized SILAC ratios were log_2_ transformed and plotted with Graphpad Prism. Phosphopeptides were classified as increased or decreased when log_2_(ratio) > 0.5 or log_2_(ratio) < -0.5, respectively, in both forward and reverse experiments. Pathway analysis was performed using Ingenuity Pathway Analysis (Qiagen). Amino acid probabilities within increased and decreased phosphopeptides were calculated using weblogo.threeplusone.com. Phosphomotif enrichment within both groups was performed using phosphosite.org/staticMotifAnalysis (algorithm: MotifAll, background selection: automatic). Identification and quantification of phosphorylated P-x-S/T-P sites was performed using a custom-made Python (v3.6) script that screens all phosphopeptides for the presence of said motifs in which Maxquant indicated the threonine/serine to have a phospho (STY) probability > 0. The raw mass spectrometry proteomics data of the strain-regulated phosphoproteome have been deposited to the ProteomeXchange Consortium via the PRIDE (*59*) partner repository with the dataset identifier PXD052926.

### RNA sequencing

Technical quadruplets were randomly divided over seperate 6-well stretch devices to prevent potential positional bias. After 2 hours of strain application, cell lysis and RNA isolation were performed using the NucleoSpin RNA mini kit for RNA purification (Macherey-Nagel, 740955.250) with DNase treatment. cDNA was synthesized using the iScript cDNA synthesis kit (Bio-Rad, 1708891). The quality and quantity of the RNA was measured using NanoDrop 2000 spectrophotometer (Thermo Scientific). Sequencing was performed using an Illumina NextSeq2000 (1 x 50 bp). Quality control on the sequence reads from the raw FASTQ files was done with FastQC (v0.11.8). TrimGalore (v0.6.5) was used to trim reads based on quality and adapter presence after which FastQC was again used to check the resulting quality. rRNA reads were filtered out using SortMeRNA (v4.3.3) after which the resulting reads were aligned to the reference genome CanFam4_GSD_1.0 using the STAR (v2.7.3a) aligner. Follow-up quality control on the mapped (bam) files was done using Sambamba (v0.7.0), RSeQC (v3.0.1) and PreSeq (v2.0.3). Readcounts were then generated using the Subread FeatureCounts module (v2.0.0). Filtering of lowly expressed genes (reads > 10 in at least two samples were included), normalization for composition bias, log differentiation and differential expression analysis (stretched vs. unstretched samples) was performed with R packages EdgeR (v4.3.0) and Limma (v4.3.0) using the limma-voom pipeline. Differentially expressed genes were defined as those with fold-change of at least 1.5 and *p* value of less than 0.05. R package EnhancedVolcano (v4.3.0) was used to generate the volcanoplot. Pathway analyses were performed using Ingenuity Pathway Analysis (Qiagen). The raw transcriptomics data of the strain-induced gene expression changes have been deposited to the Gene Expression Omnibus (GEO) with the dataset identifier GSE290599.

### Western blotting

Following stretch or EGF (Peprotech, AF-100-15, 60 nM) treatment, cells were lysed directly with Laemmli sample buffer. Proteins were denatured for 10 minutes at 95°C, separated by SDS-PAGE, and transferred to PVDF Immobilon Transfer Membranes (Merck) by Western blotting. Membranes were blocked with 2% BSA in Tris-Buffered Saline for 30 minutes at 4°C. Proteins of interest were probed by overnight incubation with primary antibodies at 4°C. pERK (Cell Signaling Technology, 4370), ERK (Cell Signaling Technology, 4370), and pMEK (Cell Signaling Technology, 9121) antibodies were used 1/2500. pEGFR (Cell Signaling Technology, 2234), EGFR (Cell Signaling Technology, 4267) and MEK (Cell Signaling Technology, 9122) antibodies were used 1/1000. α-Tubulin (Sigma, T9026) and p130Cas (BD Biosciences, 610272) antibodies were used 1/5000. Primary antibody detection was done by 1 hour incubation with either Horse Radish Peroxidase-(Dako) or Alexa-(Invitrogen) conjugated secondary antibodies at 4°C. HRP-incubated membranes were imaged upon Enhanced Chemiluminescence using ImageQuant LAS 4000 mini (GE Healthcare Life Sciences), Alexa-incubated membranes were imaged directly using Typhoon Biomolecular Imager (Amersham). Western blots were quantified using the Gel Analysis tool in ImageJ.

### Proximity labeling

APEX2-expressing cell lines were plated on 12 mm glass coverslips coated with 50 μg/mL Rat Tail Collagen Type 1 (Corning) for immunofluorescence experiments or in 15 cm tissue culture dishes (2 per condition) for mass-spectrometry experiments. 24 hours after plating, cells were incubated with 500 µM biotin-phenol in growth medium for 30 minutes at 37°C. Biotinylation was initiated by adding H_2_O_2_ to a final concentration of 1 mM for 1 minute at room temperature. The medium was removed, and cells were washed three times with quencher solution (10 mM Trolox-500, 5 mM L-ascorbic acid, and 5 mM sodium azide (NaN_3_) in DPBS (Lonza)) to quench the biotinylation and to reduce post-fixation/post-lysis inclusion of biotin-phenol.

For immunofluorescence experiments, cells were fixed in 4% (v/v) paraformaldehyde in DPBS for 20 minutes at room temperature, permeabilized with Triton X-100 for 3 minutes at room temperature and blocked with IF blocking solution (2% Bovine Serum Albumine (BSA), 3% Normal Goat Serum (NGS), and 50 mM ammonium chloride (NH_4_Cl) in DPBS) for 1 hour at room temperature. Biotinylated proteins were stained with Alexa Fluor 568-conjugated Streptavidin (1/500, Fisher Scientific, 10297422) for 1 hour at room temperature. Imaging was performed using an inverted Zeiss Cell Observer widefield microscope with 20x dry objective (Fluar, N.A. 0.75), Images were captured using ZEN software (Zeiss, ZEN v.2.3) and processed with ImageJ.

For mass-spectrometry experiments, cells were harvested on ice in ice cold RIPA lysis buffer (1% (w/v) Triton X-100, 0.5% (w/v) Sodium Deoxycholate (NaDOC), 0.05% (w/v) sodium dodecyl sulfate (SDS), 50 mM Tris-HCl, pH 7.5, 150 mM sodium chloride (NaCl), and 200 µM EDTA, supplemented with quenchers, protease inhibitors Aprotinin and Leupeptin, and phosphatase inhibitors (NH_4_VO_3_, NaF). Lysates were then cleared by centrifugation at 15,000 RPM for 10 minutes at 4°C. Remaining biotin-phenol was filtered out of the lysates by three sequential dialysis steps (2x 3 hours, 1x overnight) at 4°C in ice cold RIPA lysis buffer supplemented with protease and phosphatase inhibitors using a 3.5K MWCO Spectra/Por Dialysis Membrane (Spectrum Laboratories). For pull-down of biotinylated proteins, dialysed lysates were incubated with pre-washed streptavidin-agarose beads (high capacity: >10 mg biotinylated BSA/mL; Thermo Fisher Scientific) for 1 hour at 4°C. Streptavidin beads were washed by five sequential washing steps to remove nonspecific binders and SDS: twice with ice cold RIPA buffer, once with 1 M potassium chloride, once with 0.1 M sodium carbonate, and once with 2 M urea in 10 mM Tris-HCl (pH 8.0). Beads were mixed with 8 M urea (MP Biomedicals) in 1 M ammonium bicarbonate (ABC) (Fluka) and diluted 4x to 2 M ureum. Cysteine alkylation was induced by adding 2 chloroacetamide (CAA) (Sigma) to the samples. On bead tryptic digestion was performed by incubating diluted samples overnight with 200 ng Trypsin/Lyc-C (Promega) in 50 mM Tris-HCl, pH 8, at 37°C. Peptides were desalted by loading the samples onto an in house C18 stagetip, washed with FA and eluted in 80% ACN and FA. Isolated peptides were separated by inline liquid chromatography (LC) on a 30 cm column (50 µm ID (New Objective)) packed with 3 µm aquapur gold C-18 material (dr. Maisch) using 140 minute gradient (7% to 80% ACN, 0.1% FA), and peptides were electro-sprayed into an Orbitrap Fusion Tribrid Mass Spectrometer (Thermo Fisher Scientific). Peptides were detected by a MS1 precursor scan (m/z: 400-1,350) followed by a Top Speed MS2 scan in a data-dependent mode, with the resolution over the full-scan set to 240,000 and the intensity threshold at 5,000 ions. Most intense ions were isolated by the quadrupole and fragmented with an HCD collision energy of 35%. The maximum injection time of the iontrap was set to 100 milliseconds. Raw data was analyzed with MaxQuant software (v1.5.2.8) (*58*) using the iBAQ algorithm. Methylation was set as fixed modification and methionine oxidation as variable modification. For peptide identification, the *canis lupus familiaris* Uniprot FASTA database was searched with protein false discovery rate set to 1%. Proteins identified with two or more unique peptides were filtered for reverse hits and standard contaminants. Scatter plots were created with Perseus (v1.5.5.3). The raw mass spectrometry proteomics data of the proximity labeling have been deposited to the ProteomeXchange Consortium via the PRIDE (*59*) partner repository with the dataset identifier PXD061223.

### ADAM enzymatic activity measurements

Following 15 minutes application of stretch, cells were fixed in 4% (v/v) paraformaldehyde in DPBS for 20 minutes at room temperature and washed 2x with DPBS. PDMS membranes were punched out and mounted between glass coverslips using Immuno-Mount (Thermo Fisher Scientific) and Z-stacked images (2.1 um slices with approximately 15% overlap) were obtained using an inverted Zeiss LSM880 confocal Laser Scanning Microscope with a 40x oil immersion objective (Plan-Neofluar, N.A. 1.30). Single junction ROIs were drawn manually over individual cell-cell junctions in max projections of the mNeon channel in ImageJ (NIH) using 5 pixel wide freehand line segments. To determine mScarlet/mNeon ratios of individual cell-cell junctions, an R script was developed that identifies the optimal Z-slice of each individual ROI based on mScarlet intensity and calculates the average intensity ratios within that particular Z-slice for each ROI. Experiments that investigated the effect of a perturbation (TAPI-2 or T151-E-cadherin) on strain-induced ADAM activity were included in analysis if mechanical strain decreased mScarlet/mNeon ratio > 10% in the control (untreated or wild type, respectively). Ratio plots were made using Graphpad Prism.

## Acknowledgements

We thank members of our laboratories for helpful discussions.

## Author contributions

R.M.H., M.C.v.d.N., W.J.P. and M.G. wrote the manuscript and assembled the figures and tables. R.M.H., M.C.v.d.N., H.R.V., B.L.P., M.M., W.J.P. and M.G. designed the experiments and contributed additional edits. W.J.P. and M.G. had the overall study responsibility and directed the research. R.M.H., M.C.v.d.N., M.J.V., T.E.J.C.N., L.v.U. and W.J.P. performed the experiments. R.M.v.E. and H.R.V. performed mass-spectrometry measurements and analysis. J.Y.S., M.A.H. and B.L.P. designed and created the 6-well cell strain device and its controller setup. E.D., K.T. and M.M. designed and created the EREG and NECL5 shedding reporters.

## Data and materials availability

The raw mass spectrometry proteomics data have been deposited to the ProteomeXchange Consortium via the PRIDE partner repository with the dataset identifiers PXD052926 (strain-regulated phosphoproteome) and PXD061223 (E-cadherin proximity labeling). The raw transcriptomics data (strain-induced gene expression changes) have been deposited to the Gene Expression Omnibus (GEO) with the dataset identifier GSE290599.

