## Supplemental Figures for "E-cadherin mechanotransduction activates EGFR-ERK signaling in epithelial cells by inducing ADAM-mediated ligand shedding"

**Supplementary Materials**

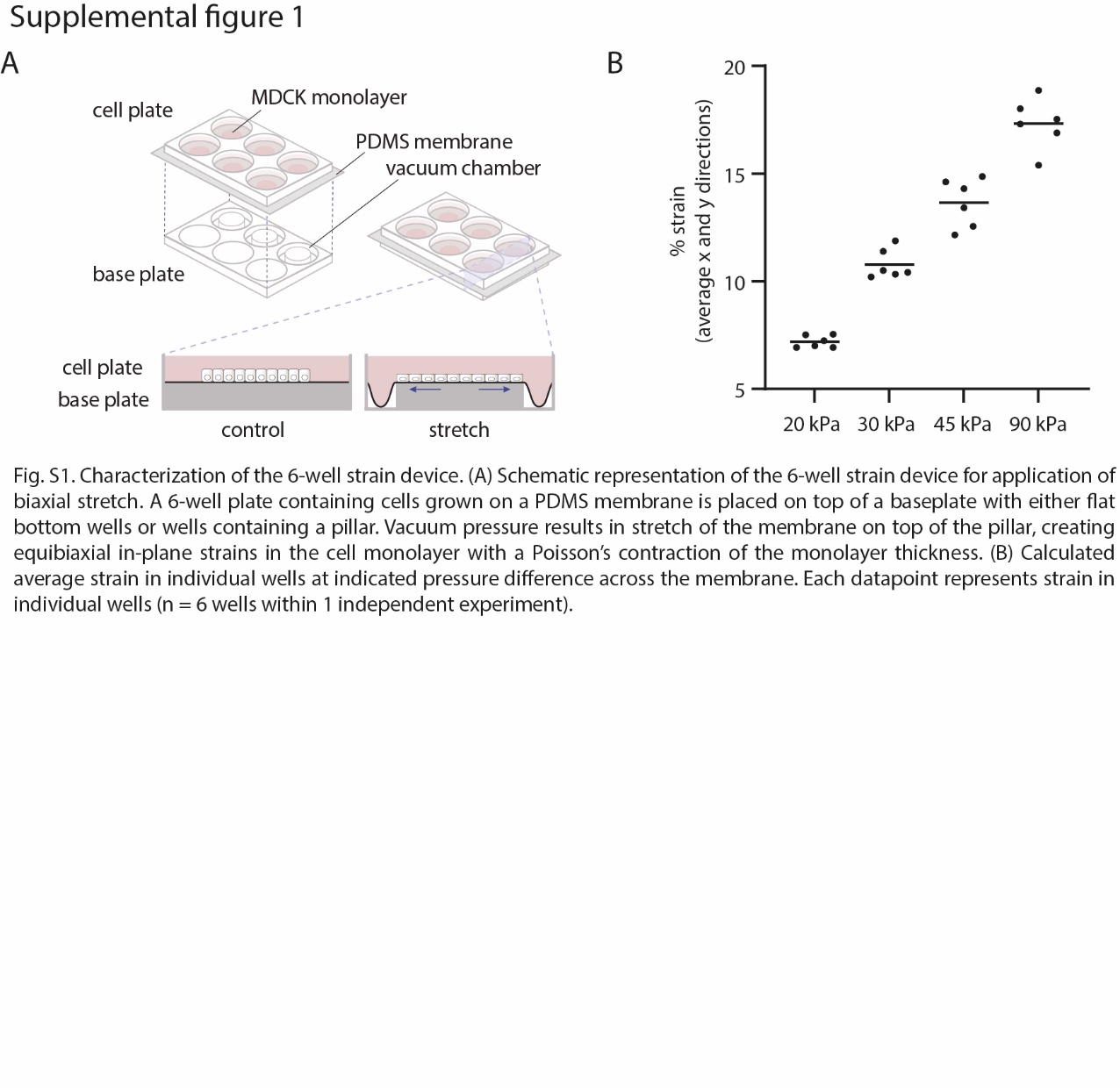

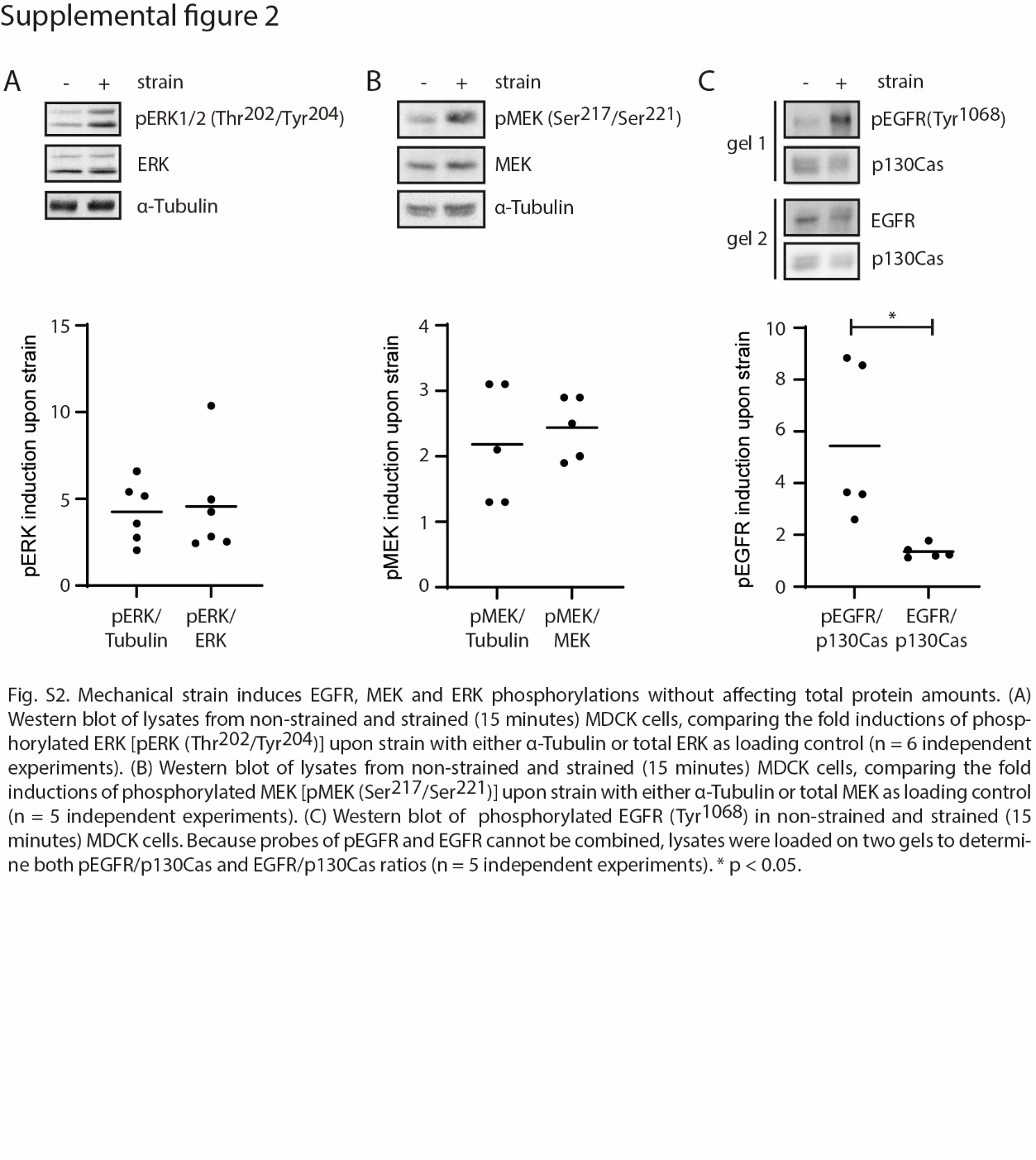

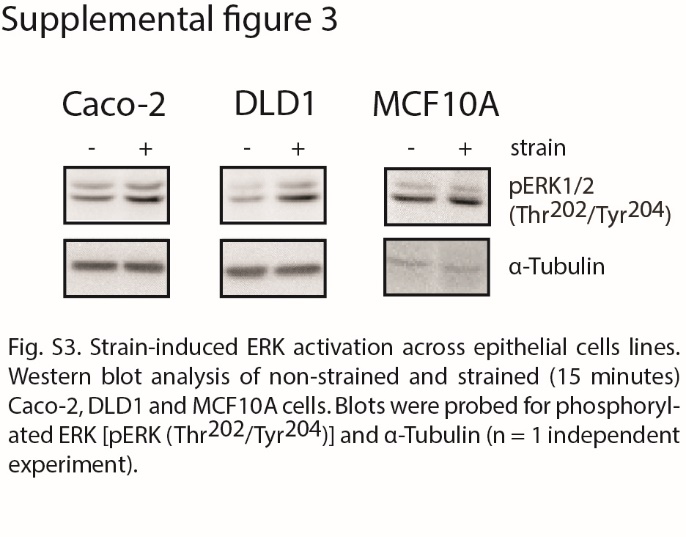

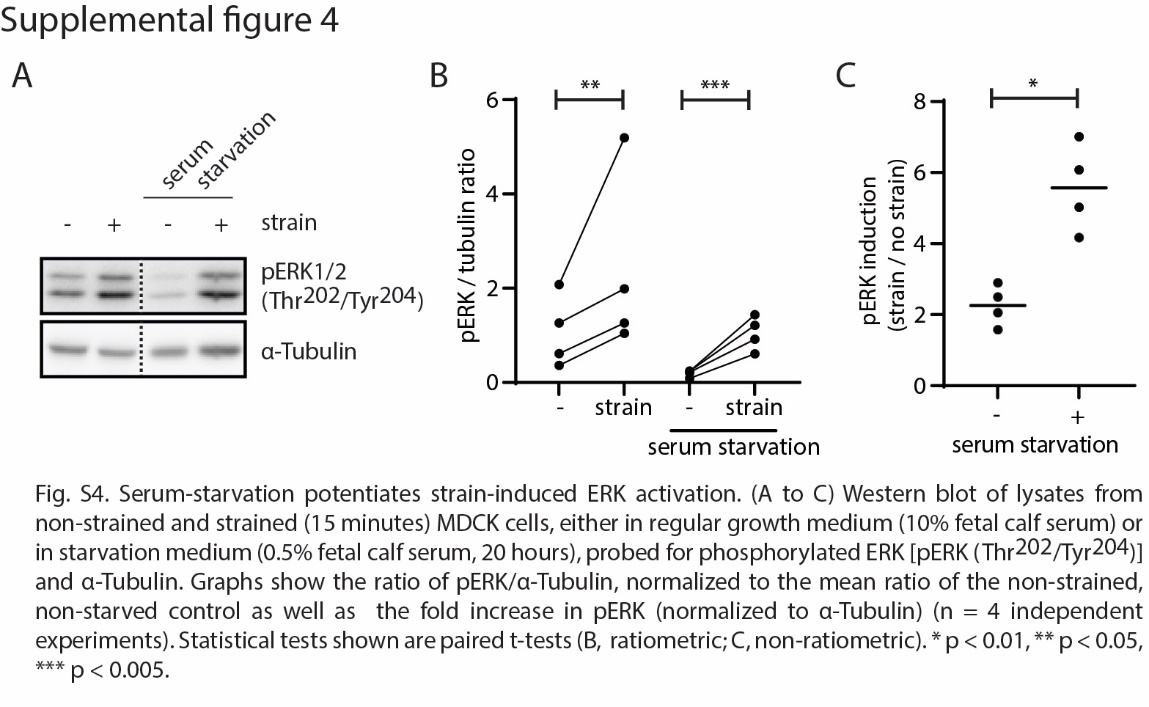

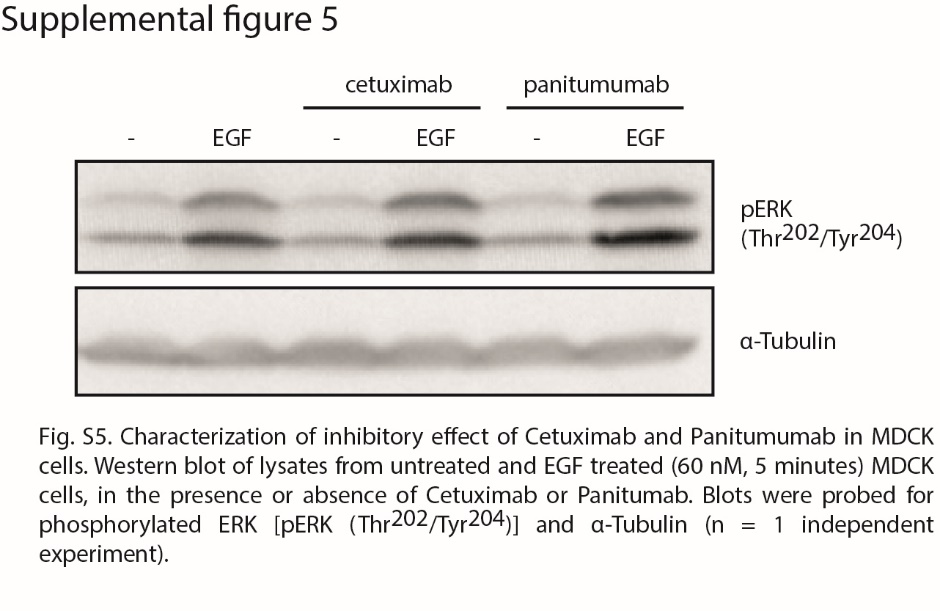

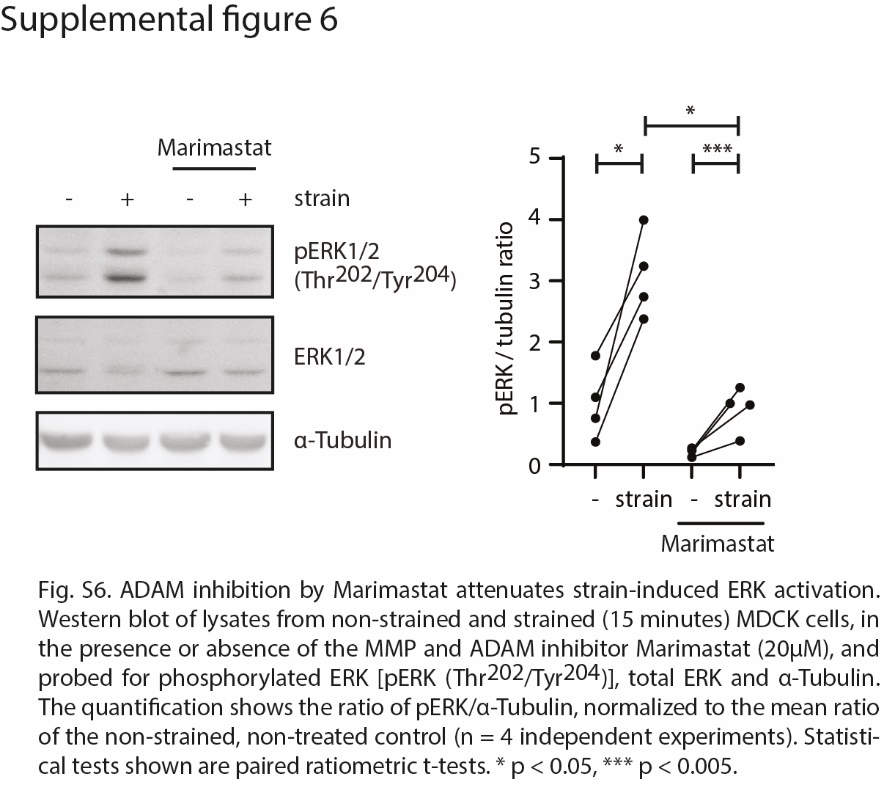

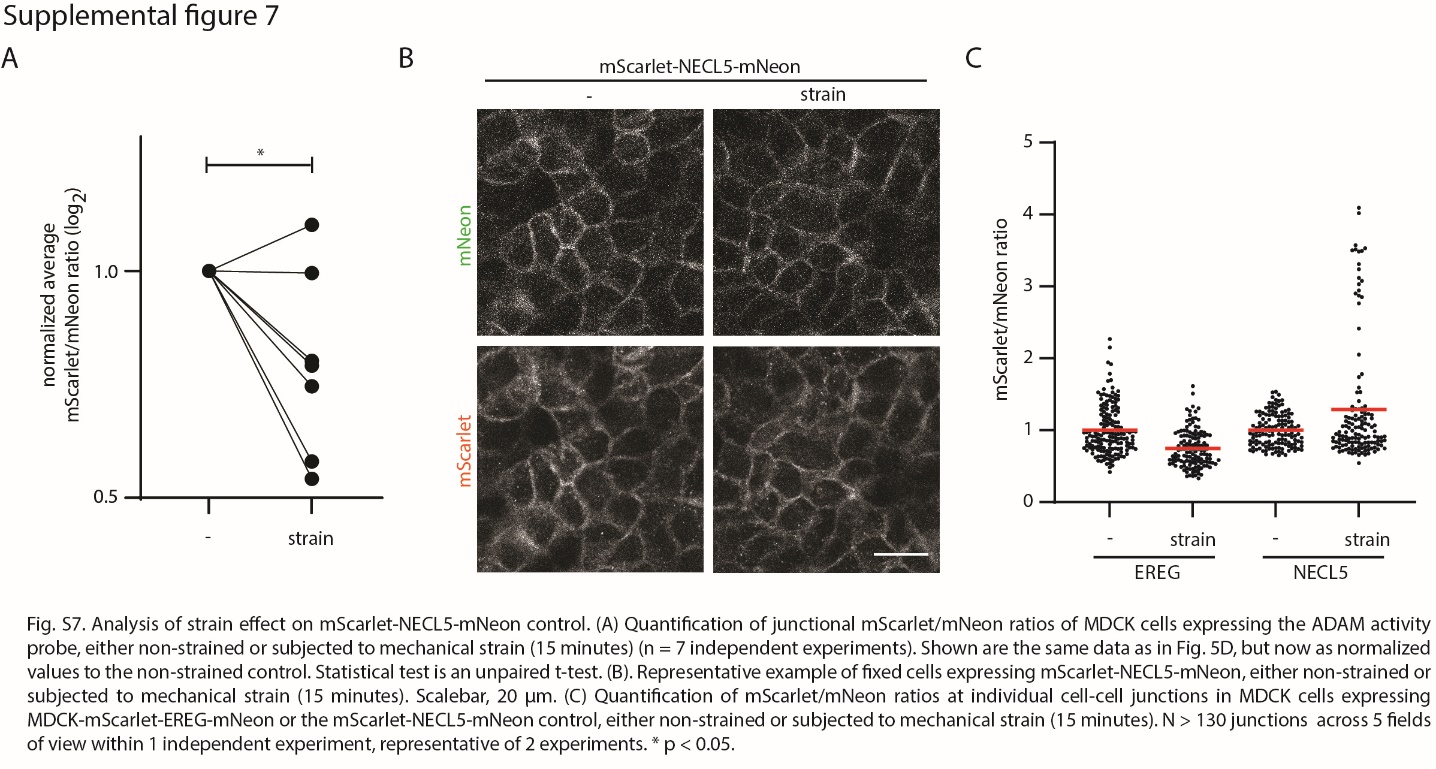

**List of supplementary data files**

Data File S1: Phosphoproteomics of strained and non-strained MDCK cells.

Data File S2: Ingenuity Pathway Analysis (IPA) of strain-regulated phosphorylated proteins.

Data File S3: RNA sequencing of strained and non-strained MDCK cells.

Data File S4: Literature based selection of transcriptional targets of ERK signaling.

Data File S5: Mass spectrometry measurement of E-cadherin-APEX2 and APEX2-NLS MDCK cells.
